# Using big data to estimate whale watching effort

**DOI:** 10.1101/2020.11.30.403923

**Authors:** Javier Almunia, Patricia Delponti, Fernando Rosa

**Affiliations:** Loro Parque Fundación, Puerto de la Cruz, Spain; Aula de Comunicación, Departamento de Ciencias de la Comunicación y Trabajo Social, Facultad de Ciencias Sociales y de la Comunicación, Universidad de La Laguna, La Laguna, Spain; Laboratorio de Bioacústica Física y multi Sensores Distribuidos, Departamento de Ingeniería Industrial, Universidad de La Laguna, La Laguna, Spain

**Keywords:** automatic identification system, cetacean, whale watching, carrying capacity, sustainability

## Abstract

The growing concerns about the negative effects caused by whale watching on wild The growing concerns about the negative effects caused by whale watching on wild cetacean populations are evincing the need to measure whale watching effort more precisely. The current alternatives do not provide sufficient information or imply time-consuming and staff intensive tasks that limit their effectiveness to establish the maximum carrying capacity for this touristic activity. A methodology based on big data analysis, using Automatic Information System (AIS) messages, can provide valuable vessel activity information, which is necessary to estimate whale watching effort on cetacean populations. We used AIS data to automatically detect whale watching operations and quantify whale watching effort with high spatial and temporal resolution in the Canary Islands. The results obtained in this study are very encouraging, proving that the methodology is able to estimate seasonal and annual trends in the whale watching effort. The methodology has also proved to be effective in providing detailed spatial information about the whale watching effort, which makes an interesting tool to manage spatial regulations and enforce exclusion zones. The widespread use of AIS devices in maritime navigation provides an enormous potential to easily extend this methodology to other regions worldwide. Any public strategy aimed at the sustainable use of the marine resources should enhance the use of this kind of information technologies, collecting and archiving detailed information on the activity of all the vessels, especially in marine protected areas.

## 1 INTRODUCTION

There is an increasing number of people that are demanding whale watching boat trips worldwide, fueling a fast growing industry that already accounted for 3,300 operators by the end of the previous decade (O’Connor et al., 2009). As this activity is not based on a lethal or consumptive use of the cetaceans, whale watching has been often labeled as “green”, “eco-friendly” or “sustainable” tourism (Schuler et al., 2019). However, early in this century the first evidence about short-term behavioural changes provoked by vessel density appeared (Allen and Read, 2000) and since then many authors have reported negative impacts of whale watching activities in different cetacean species (Erbe, 2002; Lusseau, 2003; Lusseau and Higham, 2004; Constantine et al., 2004; Orams, 2004; Lusseau et al., 2006, 2009; Schaffar et al., 2009; Parsons, 2012; Christiansen et al., 2014). These short-term behavioural changes included: surfacing/diving, agonistic behaviour, anti predator behaviour, acoustic, group size or cohesion, swimming speed, swimming direction, altered feeding or resting, and altered respiratory frequency (see Parsons (2012) for a complete review). Shortly after, the first evidence of the long-term negative impacts produced by whale watching appeared in one of the best-studied dolphin populations (Bejder et al., 2006), confirming the concern of the International Whaling Commission (IWC), who in 1997 created a working-group to monitor whale-watching sustainability (IWC, 2004). In that sense whale watching, like most other human activities, can be considered an evolutionary selection force, which alters the life of the targeted population (Lusseau et al., 2006). The fact that cetacean (whale, dolphin, and porpoise) watching is the greatest business reliant upon cetaceans worldwide (Parsons, 2012), targeting at least 56 (including endangered and threatened) species in all oceans so far (Bejder et al., 2006), urges to find sustainable ways to perform these activities.

To accomplish this goal, it is essential to determine the carrying capacity, or maximum whale watching effort, that any cetacean population can bear in a sustainable way. The need to evaluate carrying capacity in whale watching activities has been early identified in the scientific literature (Curtin, 2003; Higham et al., 2008; Andreu et al., 2009) but the intensity or effort of the activity has been rarely considered as a factor in the impact studies. Traditionally the whale whatching effort has been assumed to be proportional to the number of vessels operating in a certain area. But this is a deficient measurement, as it does not consider the different activity budgets of each vessel, its physical characteristics, or the seasonal and geographical variations on the whale watching events. More recently, some studies have used land-based visual observations (with binoculars or theodolite) and also acoustic data in order to measure whale watching intensity (Pirotta et al., 2015; Schuler et al., 2019), determining the concurrent number of vessels, or the total time spent in the proximity of the animals. This methodology is much more precise and appropriate to establish the effect of different whale watching intensities in the short-term behavioural disturbances produced on the cetaceans. But it is also geographically limited, enormously time consuming and staff intensive, which makes its application on regional monitoring programs quite unrealistic. Similarly, the need to obtain precise effort measures has been also highlighted by the authors of the mathematical models proposed to address the long-term sustainability of tourist interactions with cetaceans (Higham et al., 2008; Lusseau et al., 2009; New et al., 2020), as the quality of the model projections will heavily depend on the amount and quality of the whale watching effort data available.

The Canary Islands is one of the top whale watching destinations worldwide. In 1998 Spain was considered among the three countries that could claim to have taken over one million people whale watching in one year (Connor 2008) mainly thanks to the visitors registered in the Canaries. Ten years after, despite a visitor reduction due to regulatory measures and weather issues, the Canary Islands were considered the fourth whale watching destination worldwide with 611.500 whale watchers per year (Connor 2008). Whale watching in the Canaries is strongly focused on Tenerife Island, which accounts for an estimated 85% of total whale watchers (Connor 2008), around a resident population of some 350-450 short-finned pilot whales *(Globicephala macrorhynchus)* along with transient visitors can be found off the south-west coast of Tenerife (Canary Islands) mainly in water depths from 800 to 2000 m (Heimlich-Boran and Heimlich-Boran, 1990; Aguilar Soto et al., 2008). The fact that the Canary Islands is one of the leading whale watching destinations worldwide, and the concentration of this activity in a well defined resident species, constitutes an ideal laboratory to study whale watching effort.

The Automatic Identification System (AIS) is an unattended vessel reporting system that was developed for collision avoidance. The AIS transponder automatically broadcasts information on a vessel’s name, position, course, speed, etc. at regular intervals to all AIS receivers in the area. Due to an International Maritime Organization (IMO) mandate, many vessels are now directed to carry AIS systems and broadcast AIS messages (Lapinski and Isenor, 2011). Originally designed for radar augmentation and vessel traffic services (VTS), the system can be also used to collect information about traffic in the area with little effort. AIS provides position updates at sample rates varying from three seconds to three minutes dependent on the individual vessel’s manoeuvre situation (Aarsæther and Moan, 2009). Apart from its original goal, the enormous AIS data available has proved to be a valuable source of information on human use of marine areas. As a consequence, it has been used for different purposes; from monitoring fishing activity and protected area regulation compliance (Natale et al., 2015; de Souza et al., 2016; Rowlands et al., 2019), to evaluate cetacean-vessel collision risks (Greig et al., 2020), but never before to evaluate whale watching effort. The aim of this study is to evaluate the potential use of AIS data, combined with an open source Digital Terrain Model (DTM), to automatically measure the whale watching effort on a specific region.

## 2 METHODOLOGY

### 2.1 AIS data

Two specific geographical areas were selected to characterize the whale watching activities off Southern Tenerife island (27.9-28.4°N; 16.5-17.0°W) and off Southern Gran Canaria island (27.5-28.0°N; 15.5-16.0°W). The vessels included in this study were selected searching, at the MarineTraffic© database, for the names of the ships authorized by the Canary Islands Government to perform whale watching activities in the region. The MMSI (Maritime Mobile Service Identity) numbers of all the authorized ships that were equipped with an AIS transponder, and consequently appeared in the MarineTraffic© database searches, were included in the study. The search resulted in a total of 23 vessels (out of 120 authorized for whale watching activities in the region) that produced AIS messages between 2016 and 2020.

AIS data were collected by Exmile Solutions Ltd. (London, UK) from the calendar years 2016 to 2019 and part of 2020, and received either from terrestrial stations or satellite. To ensure effective management of the incoming information in the database, MarineTraffic© uses proprietary down-sampling techniques not to archive consecutive positions within minutes. This results in a maximum resolution of one minute for the archived data. All the available archived AIS messages, received from vessels selected for the study, were obtained from the database MarineTraffic©. The available messages were previously filtered by time stamp between ‘2016-01-01 00:00’ and ‘2020-03-14 00:00’, and further filtered to select just the operational hours of the whale watching vessels (between 09:30 and 17:30).

The original AIS data received had 729,951 messages (10,558 of them from satellite and the rest from terrestrial stations). The data was not evenly distributed over the years, but showed an increasing trend related with a growing number of boats equipped with AIS transponders in the region: 2016 (72,751), 2017 (135,158), 2018 (217,457), 2019 (261,619) and 2020 (42,966 just in two and a half months). The data was processed using scripts written by the authors using the programming language Python (Van Rossum, Guido and Drake, 2009) running in Anaconda Spyder. A preliminary data quality control was performed removing all the positions out of the geographical range of the study.

### 2.2 EMODnet Digital Terrain Model

As the original AIS messages do not include information about the depth at the vessel’s location, it was extracted from a Digital Terrain Model (DTM). The “EMODnet Digital Bathymetry (DTM)” is a multilayer bathymetric product for Europe’s sea basins. The DTM is based upon more than 7,700 bathymetric survey data sets and Composite DTMs that have been gathered from 27 data providers from 18 European countries and involving 169 data originators (Consortium, 2016; Thierry et al., 2019). The gathered survey data sets can be discovered and requested for access through the Common Data Index (CDI). The depth at vessel’s location was estimated for every AIS message, and all the messages that were broadcasted from positions shallower than 100 m were filtered to eliminate the messages originated from harbours, moorings and coastal navigation. After eliminating the shallower positions the data set consisted in a total of 265.090 AIS messages.

## 3 RESULTS

A first analysis of the data was aimed to identify the whale watching events from the AIS information, taking in consideration that the ship speed should be reduced to follow the whales during the sighting, and that the long-finned pilot whales in the study areas are commonly distributed in water depths from 800 to 2000 m (Heimlich Boran 1993). A density plot of the AIS messages in the data set by speed and depth (Figure 1) showed a particular region between 700 and 1500 m in depth were the vessel speed was consistently under 2.5 knots. This can be considered as a clear indication of whale watching operations, specially because the figure also illustrates how the vessels typically cruise at 6 knots regardless of the depth. It is also noticeable that speeds lower than 2.5 knots couldn’t be find in those water depths where the whales were absent.

**Figure 1.**
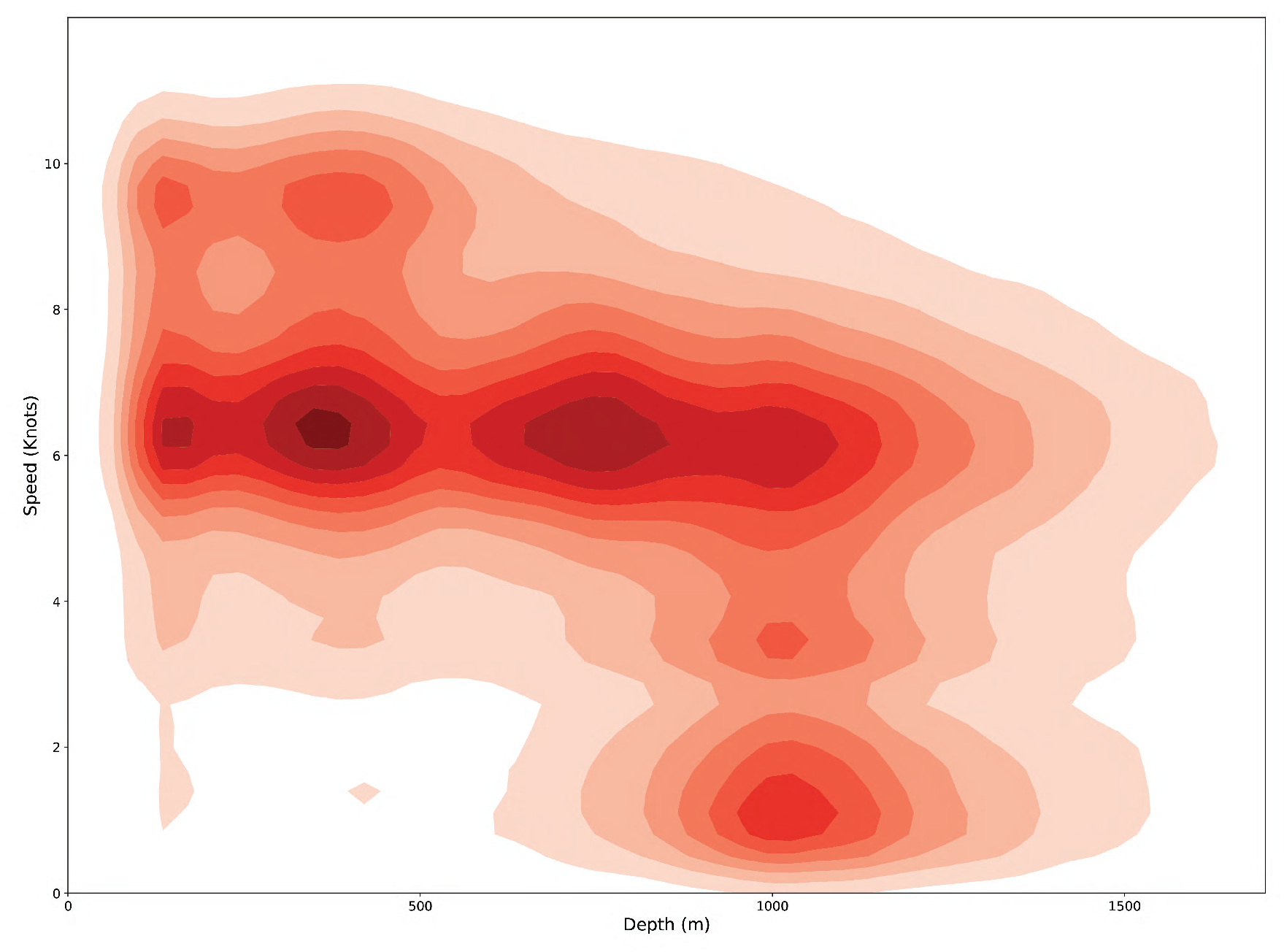
Density plot of the AIS messages in the data set by speed (Knots) and depth (meters)

Subsequently, all the AIS messages with speeds higher than 2.5 knots were discarded from the data set, and the frequency distribution of the remaining 43,379 AIS messages was plotted in a histogram (Figure 2). The histogram shows a clear peak around 800-1,200 m in depth, which supports the idea that the AIS messages were emitted while the vessels were performing whale watching operations with short-finned pilot whales, as this depth range matches the one described for the species in the region (Heimlich Boran 1993).

**Figure 2.**
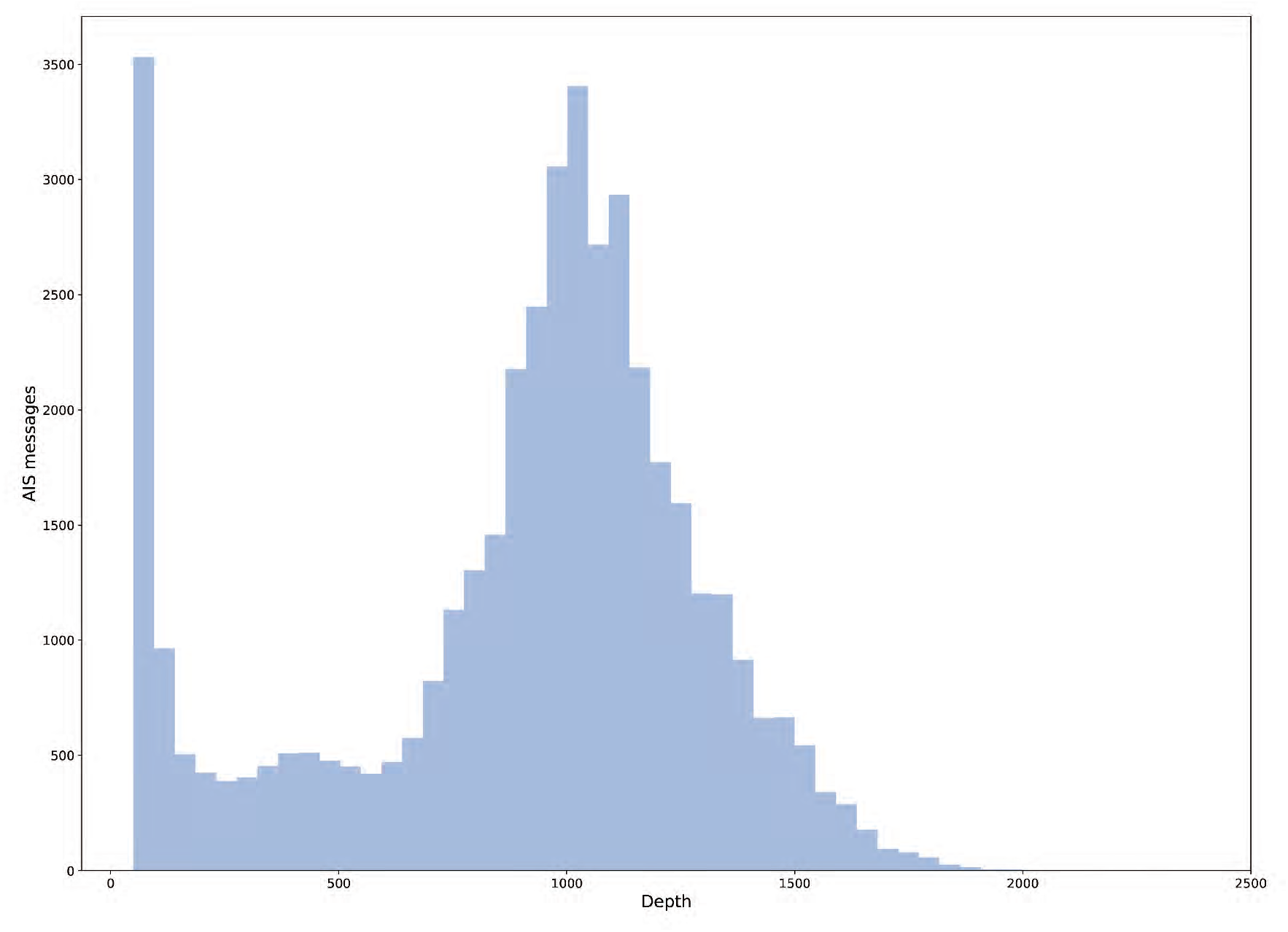
Frequency distribution of the AIS messages broadcasted at different depths in the study areas.

The geographical distribution of the density plot (Figure 3) for the whale watching events off southwestern Tenerife (blue dots) show a narrow area (aprox. 3 Km wide) that extends over 50 Km along the island slope. This area matches perfectly the distribution of the short-finned pilot whale south-west Tenerife determined by dedicated surveys (Carrillo et al., 2010). On the other hand, the density plot of the whale watching distribution off the south-west coast of Gran Canaria (Figure 4) shows a very similar structure, despite that the area is wider and more distant from shore. This difference could be expected due to the gentle island slope and much broader island platform of Gran Canaria. In both cases, the density plots indicate that whale watching events are more frequent (dark blue in the density plot) in a very small space (aprox. 1 *km*^2^) compared with the total area where the whales were present.

**Figure 3.**
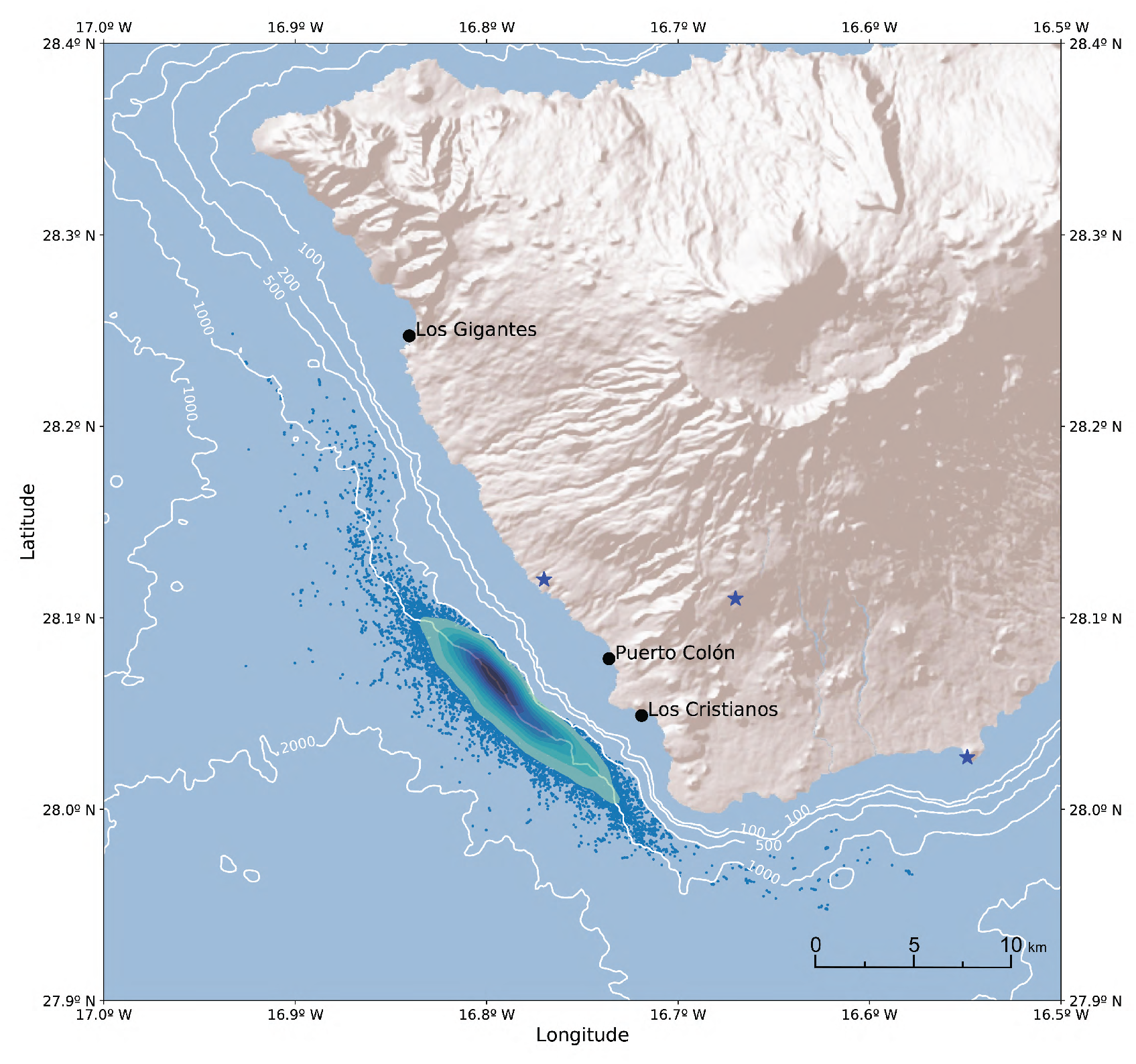
Density map of the whale watching events (small blue dots) off Tenerife Island. Black dots represent main harbors and blue stars AIS terrestrial stations.

**Figure 4.**
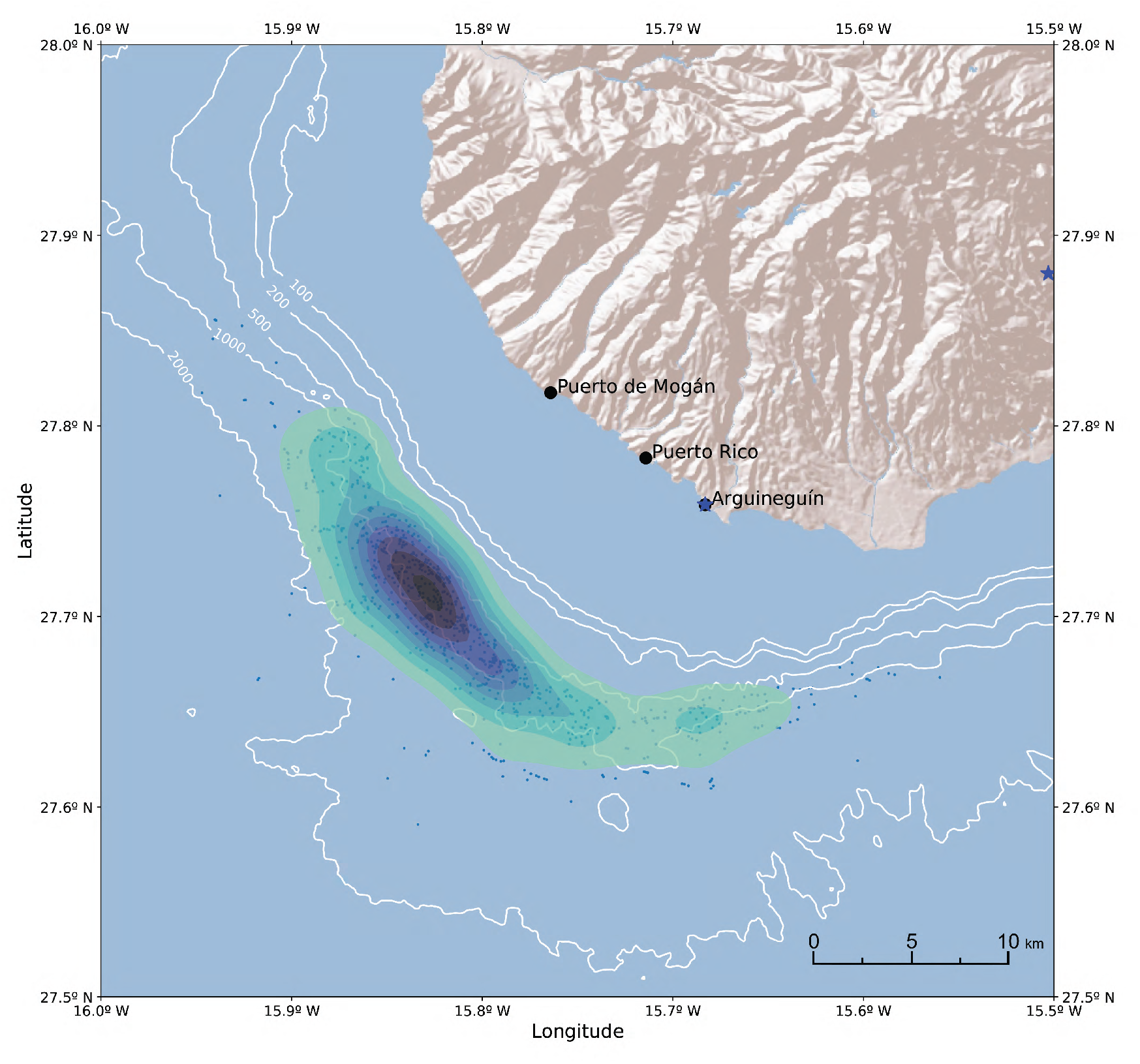
Density map of the whale watching events (small blue dots) off Gran Canaria Island. Black dots represent main harbors and blue stars AIS terrestrial stations.

To infer the duration of the whale watching events from the AIS messages, a data sequence was defined to start when the speed of the boat was under 2.5 Knots in areas deeper than 600 m. The sequence was subsequently ended when the boat reached a speed over 5 Knots regardless of the depth, suggesting that had left the whales and was cruising again. This criteria was intended to include in the same sequence several short-time ship displacements intended to approach separated animals within the same group, or to re-position the vessel while the pod was moving. As a result, a total of 8,745 sequences were identified, with a duration ranging from 1.3 to 95 minutes. The frequency distribution of the whale watching event duration (Figure 5) illustrates that the vast majority of events lasted less than 20 minutes, with few of them going over 40 min. This distribution is consistent with the actual duration of the excursions in south-west Tenerife (Pers. obs.), and falls within the maximum time allowance established in the whale watching regulations (Plasencia-Escuela, 2007).

**Figure 5.**
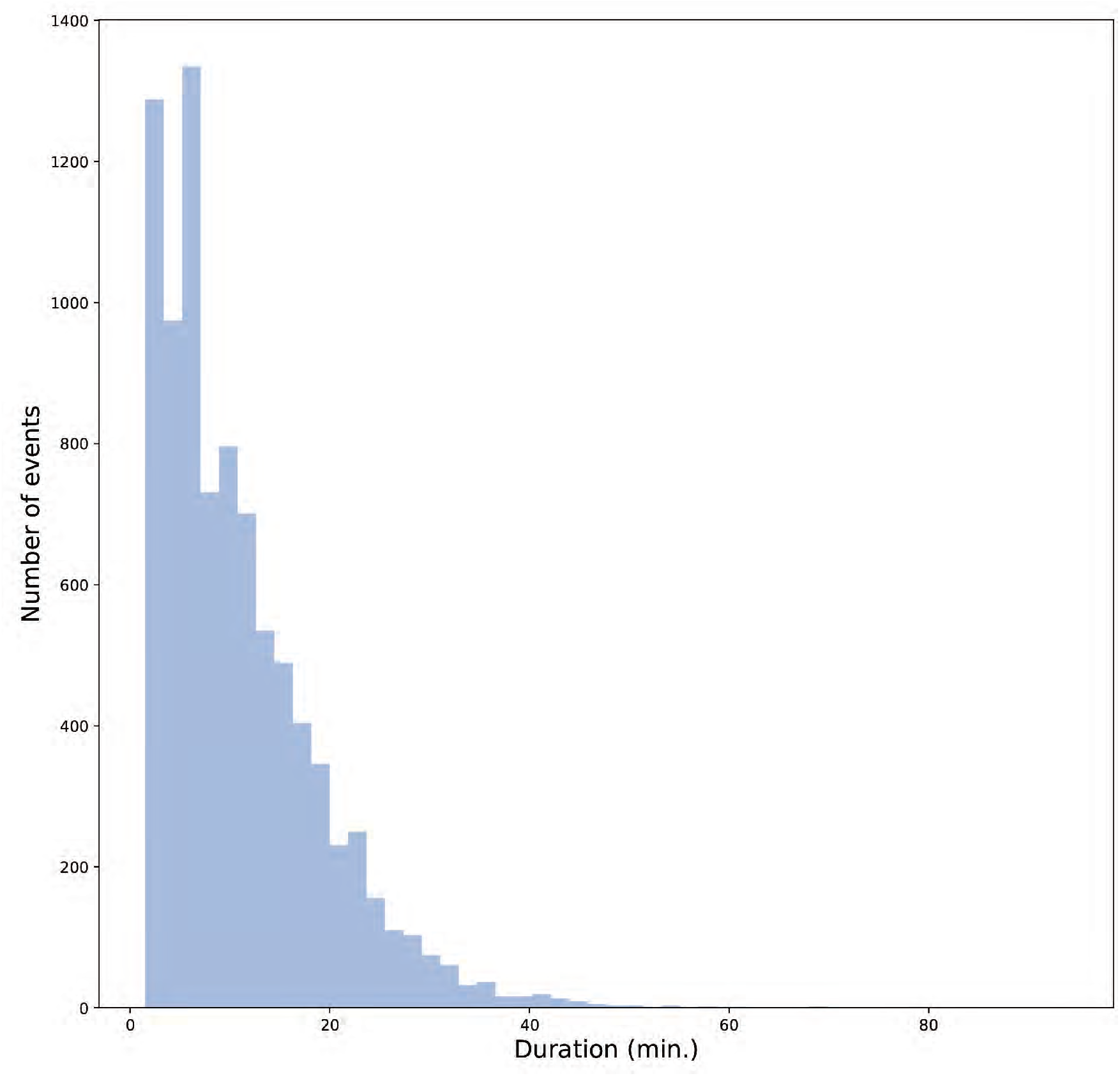
Frequency distribution of the estimated duration for the whale watching events.

Based on the inference of the duration for each whale watching event, it is possible to estimate the integrated monthly duration of whale watching by the vessels identified in the study. This parameter is an estimation of the time that the whale watching vessels were in the proximity of whales and, as a consequence, an indication of whale watching effort. The evolution of the integrated monthly duration of whale watching events during the period covered by the study seems to point out an increasing trend of roughly 3-4 fold to the end of the study. When the integrated time is normalized (divided by the number of ships that were operational on each month) the raising trend is considerably reduced (Figure 6). The operational ships were monthly estimated as any vessel that at least did one whale watching event per week, to effectively exclude those stranded for maintenance operations, operating seasonally or out of business. Taking into consideration that the number of operational whale watching vessels suffered important variations in the first half of the study, and the fact that the integrated duration of the whale watching events is more stable during the second half of the study, a clear long term trend can not be concluded from these data. Nevertheless, the graph shows a clear seasonal trend in the whale watching effort per vessel, with higher effort in summer, and much lower during winter months.

**Figure 6.**
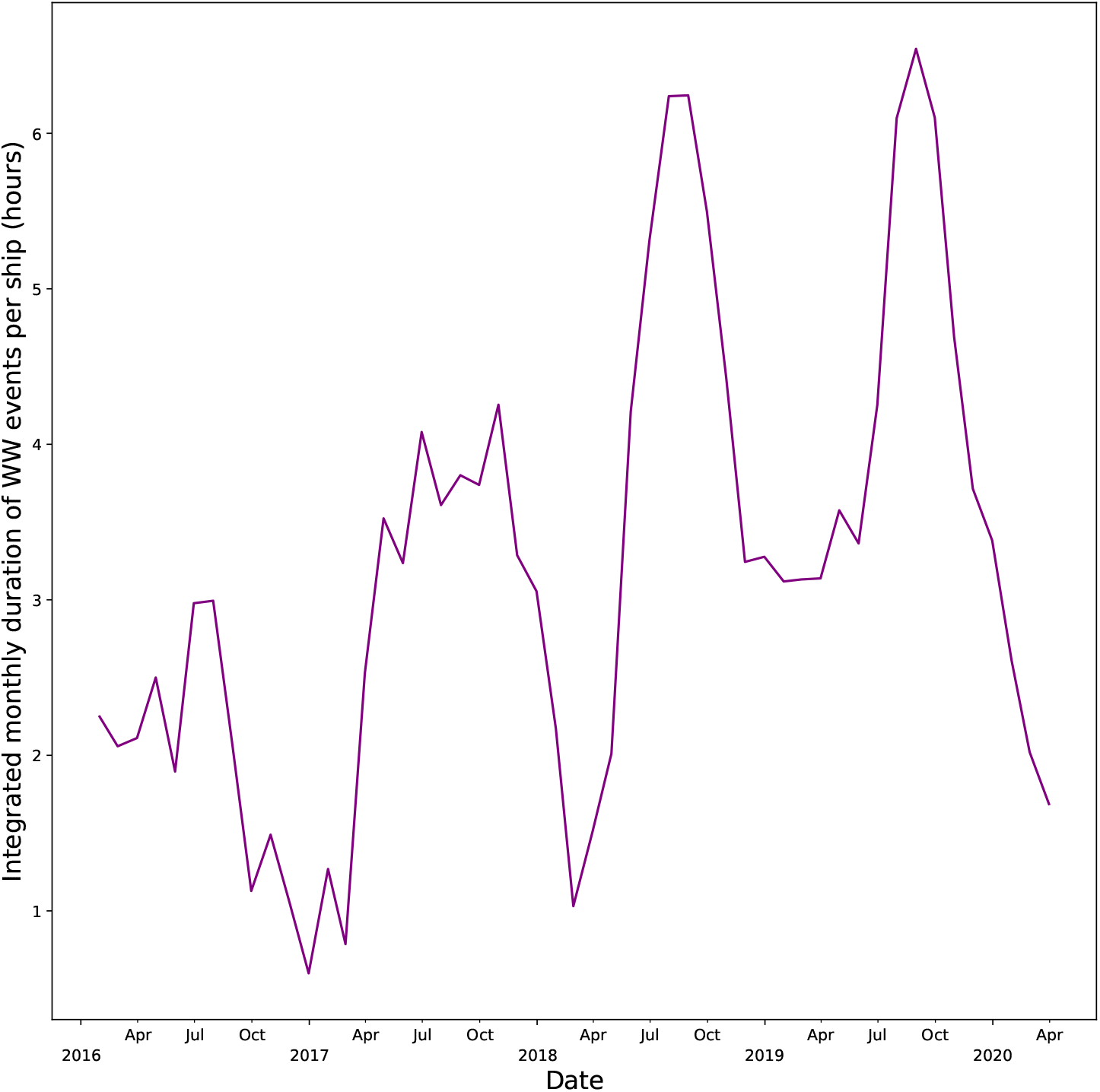
Total integrated monthly duration of whale watching events off South Tenerife normalized by the number of active vessels.

Finally, to find out if the methodology could be useful to elucidate some long term behavioural effects in the whales, like for example detecting signs of avoidance to the whale watching vessels, the AIS information was analyzed to evaluate any spatial trend in the location of the whales. As the distance of the whale watching events to shore could be biased by the low number of boats (and the fact that usually they share the daily positions by radio or simply heading to the closest whale watching boats slowly sailing in the area), the mean water depth at the whale watching events, and its latitude and longitude were used instead. Considering the extraordinary preference of this species for water depths ranging from 800 to 1,200 meters, and due to the intense island slope, if the whales were moving away from the high intensity whale watching areas, the depth associated to the AIS messages should increase. Similarly, the longitude and latitude of the whale watching events could reveal movements on the position of the animals if they were avoiding certain areas. When the daily mean depth for the whale watching events is plotted (Figure 7), there are no clear signs of avoidance trends, nevertheless a strong seasonal signal can be perceived, and appears as a significant auto-correlation of the data around a lag of 365 days. No other significant auto-correlation lags (lunar cycles or multiannual trends) could be found in the depth of whale watching events time series. Similarly, the longitude and latitude time series didn’t show auto-correlation, suggesting that the seasonal depth change of the whale watching events were the result of very subtle or inconsistent changes in the position. The seasonal depth cycle suggests that the whales are slightly closer to the island (shallower) during the summer season, and they move slightly away from the coast (deeper) during winter months.

**Figure 7.**
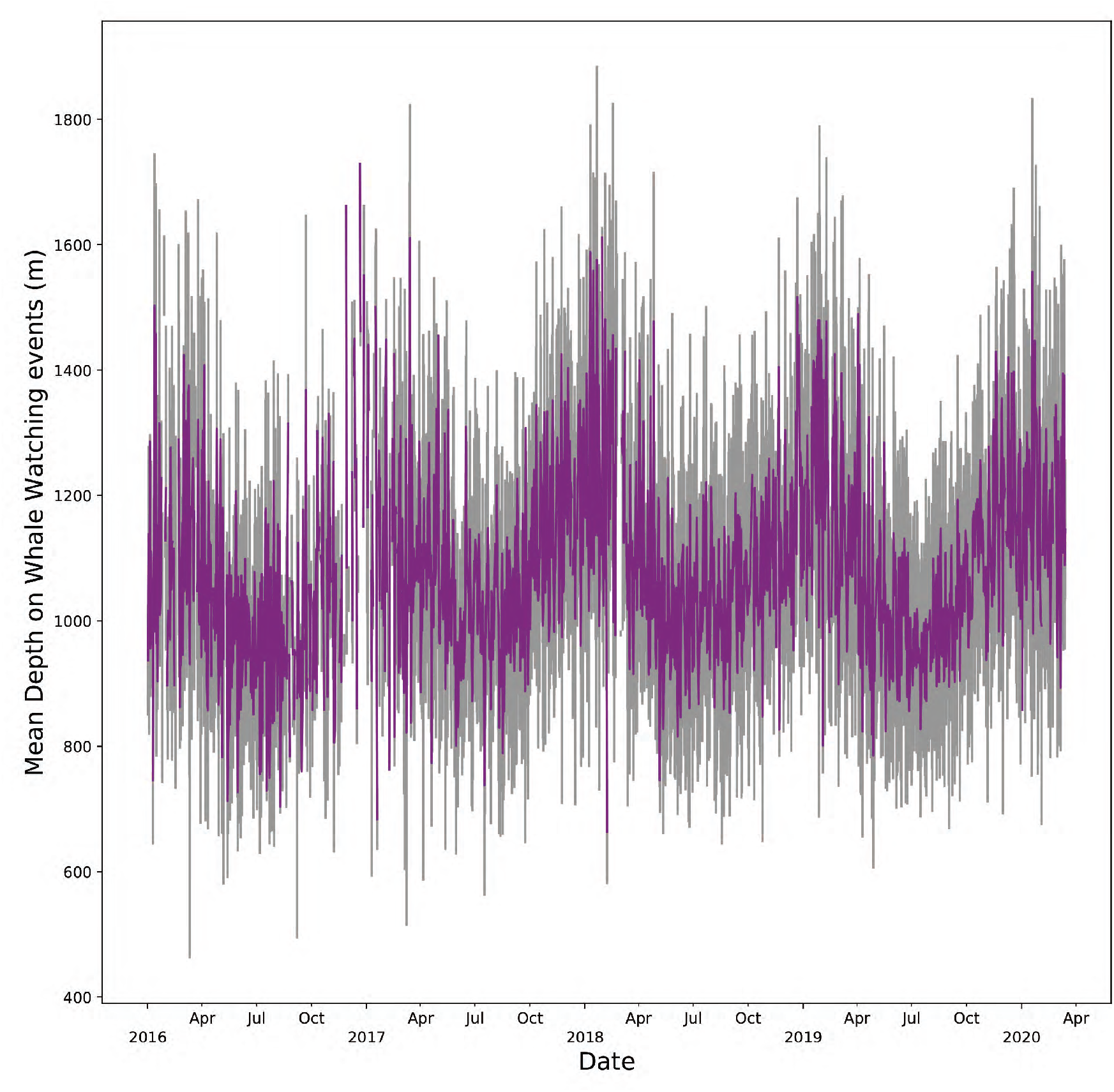
Mean daily depth of the whale watching activities off South Tenerife during the study period. Grey lines indicate standard deviation.

## 4 DISCUSSION

The fact that the vessels were not randomly chosen, and that they were not operating homogeneously during the study, does not allow to generalize the effort indexes and trends found in the results of this study. But the size of the sample (19 percent of the whale watching licensed ships in the region) gives a high confidence to evaluate the potential use of the AIS data to measure whale watching effort. The criteria established to identify whale watching events from the AIS information has proven to be solid, as the results match the distribution of the population (Carrillo et al., 2010) and the fidelity of the species to a certain bathymetric range (Heimlich-Boran and Heimlich-Boran, 1990; Aguilar Soto et al., 2008; Carrillo et al., 2010) described in the scientific literature. Notwithstanding the evidence that the criteria is valid to determine the whale watching operations with short-finned pilot whales in the Canary Islands, but it should be carefully adapted to the specific characteristics of whale watching operations focusing different species in other regions. Such a precise whale watching event measurements are scarce in the scientific literature, and usually related with direct measures (theodolite) (Schaffar et al., 2009; Cecchetti et al., 2018; Schuler et al., 2019), data collected on board (Robbins, Jooke; and Frost, 2009) or indirect estimations (ship noise)(Houghton et al., 2015). In most of the cases it is estimated just by number of licenced boats that can operate in the area, but this approach lacks of information about the amount of time that the animals were perturbated by the vessels, and also the spatial distribution of this disturbance. Obtaining detailed information about whale watching events using the theodolite method implies several observation teams of at least three people each (theodolite operator, computer operator, and 1-2 spotters)(Schuler et al., 2019). Moreover, the observations from land can be also influenced by adverse meteorological conditions (fog, dust, swell, etc.), and they are very difficult to perform in extensive or convoluted shores. In this situation, the proposed methodology is clearly advantageous as it could cover vast regions, the data collection is fully automatic and requires no staff. Actually the use of big data widens the reach of research possibilities in the Information Society. In this sense, the importance of big data, as one of the disruptive technologies in the public digital landscape, has been gradually growing, as well as the number of private organizations that in recent years have begun to store and process data to meet the demand of a market that uses and analyzes this data to generate knowledge and create business (Salvador et al., 2017). However, how to face information management, how to store it and its accessibility in the big data era, is a challenge for public endeavors, which should ensure that not only data collection is available, but also storage, interoperability and accessibility. Consequently, ensuring that AIS data from whale watching and other touristic activities is open and accessible, would imply a positive impact in sustainability.

The geographical distribution of the whale watching events off the south-west coast of Tenerife and Gran Canaria evidences the extraordinary dependence of *G. macrorhynchus* on the geomorphology of the sea bottom. It is important to highlight that this parameter has direct implications on the available optimum habitat, and it can vary dramatically in two islands within the same archipelago. The particular bathimetry of Tenerife island configures a much smaller distribution area (aprox. 150 *km*^2^ which supports 350-450 short finned pilot whales (Heimlich-Boran and Heimlich-Boran, 1990; Aguilar Soto et al., 2008). The relatively smaller size of the distribution area implies a higher risk to impact the population by an oversized whale watching industry. And, at the same time, the higher density of animals can also impose higher collision risk to the ships navigating in the area (Carrillo et al., 2010). On the other hand, the fine detail of the whale distribution obtained using AIS methodology, could be useful to define and enforce more precise low-speed areas to reduce collisions (Silveira et al., 2013; Greig et al., 2020), which has been identified as one of the measures to reduce the ship strikes in the region (Carrillo et al., 2010). In that sense, it is important to consider that the very nature of the whale sighting operations implies that each whale watching ship will preferably observe the animals closest to the shore for pure economic reasons, and that could be underestimating the density of whales in the deepest part of the range (far from coast). Similarly, the presence of whales in range areas far from touristic harbors could be underestimated for the very same reason.

The geographical coverage of AIS antennas off the south-west coast of Tenerife and Gran Canaria is obtained from the overlapping of the active stations in the area. Name, identification and location of local stations are collected in the Table 1. An appropriate field of vision of the AIS stations and their continuous operation is necessary to establish a system that automatically estimates the whale watching effort, using the proposed methodology. Both study areas have a good AIS coverage thanks to the number an position of terrestrial stations in the south of Tenerife and Gran Canaria, as proven by the low number of satellite messages in the dataset. The implementation of this methodology in other regions should be ideally based in terrestrial AIS stations that provide a good coverage of the whale watching areas and ensure a high rate of message reception. Since this navigation system is used worldwide to ensure the safety of life at sea (Tarelko, 2012), it is very likely that AIS coverage is already available, and even historical data recorded, in any region that is being used for whale watching operations.

**Table 1.** Local AIS stations relevant to the study area

| Name | Id | Latitude | Longitude |
| --- | --- | --- | --- |
| Piedra Hincada | 5659 | 28.18 | -17.66 |
| Marina San Miguel | 1067 | 28.11 | -16.67 |
| Santa Cruz de Tenerife | 2031 | 28.02 | -16.55 |
| Playa Paraiso | 2806 | 28.12 | -16.77 |
| San Sebastian | 2898 | 28.09 | -17.11 |
| Arguineguin, Gran Canaria | 5066 | 27.76 | -15.68 |
| Cruce de Arinaga | 585 | 27.88 | -15.43 |

The estimation of the whale watching event’s duration based on AIS messages seems to be quite accurate, judging by the obtained distribution of the events, which is mainly under the maximum time allowance for a sighting established by the local regulations (Plasencia-Escuela, 2007). It has to be taken into account that the highest possible precision for a whale watching sequence using MarineTraffic© archived data is two minutes, but this could be improved up to 8 seconds using a dedicated reception network. This is due to the fact that AIS transducers broadcast a message every 12 seconds when the ship is sailing at 0-14 Knots, or every 4 seconds if it is changing course (Aarsæther and Moan, 2009). A dedicated reception network would also allow the detection and storage of detailed course change events, that could be useful to estimate the duration of the whale watching events more precisely, but also to identify evasive behaviors on the whales (Schaffar et al., 2009; Christiansen et al., 2014). Despite the fact that the estimation of the sequences uses a conservative threshold to identify the end of the sighting (speed over 5 knots), it does not seem to produce a significant number of overestimated event lengths. A much more detailed study, collecting and archiving all the available AIS messages from the terrestrial stations, and comparing duration estimated by AIS with the sighting time measured by observers on board (or with theodolite from a land-based station), would be necessary to evaluate the precision of the present method. Nevertheless, even though an exact duration can not be calculated for single events, it is reasonable to assume that the errors will cancel when the values are integrated monthly. Hence, monthly integrated values could provide a good indication of the temporal trends in the whale watching intensity.

It has been proven that the behavioural disturbance in cetaceans is not only related with the presence/absence or the number of vessels in the vicinity, but also with the amount of time spent in the presence of vessels (Schuler et al., 2019). Multiple vessels simultaneously tracking a whale will accentuate this effect (Holt et al., 2009). Vessel characteristics (e.g., size and engine type) and vessel approach (e.g., angle and speed) are also likely to elicit different responses in whales (Schuler et al., 2019). Consequently, a measurement of the time that the whale watching vessels spent observing cetaceans is essential to analyse its long term consequences on the populations. The integrated duration of the monthly whale watching events calculated in this study has captured the trend of the whale watching effort better than the previous estimations. The number of licensed boats is not capable to offer seasonal differences in the whale watching activities, while those are clearly captured using the AIS messages. The maximum intensity detected in the summer season by this study matches the peak in the activity due to the higher frequency of days with better weather conditions. On the other hand, the apparent trend observed in the whale watching effort during the study could be biased by the small number of ships in the analysis and its heterogeneous activity during the whole period. To improve the accuracy of this measure, an increase in the number of whale watching vessels equipped with an AIS transponder should be necessary.

The results also show a clear seasonal trend in the average depth where the sightings were realized, with is confirmed by the impressions of some whale watching pilots, who refer that the whales tend to be closer to shore in summer, depending on weather conditions. A similar seasonal trend has been found in another well known resident population of short-finned pilot whales in Hawai’i (Mahaffy, 2000). But in that case, during the oceanographic season with the shortest day length (fall), whales were in shallower areas compared to seasons with longer days (summer) (Owen et al., 2019). Movements of odontocetes whose preys are associated with the Deep Scattering Layer (DSL) are well known (Benoit-Bird et al., 2009), and they respond to the changes in light intensity produced by lunar cycles and day-length. But the distance to the shore of Hawaiian short-finned pilot whales seems to change much more dramatically, as their dives occurred a mean of 18.3 km farther offshore (more than twice as far from shore) during a full moon compared to a new moon (Owen et al., 2019). Off the south-western coast of Tenerife the whales were consistently observed in a 3 km wide area along the coast during the length of the study (4 years), suggesting that the seasonal displacements off shore were much smaller. On the other hand, the auto-correlation of our data do not suggest the influence of lunar cycles in the position of the animals, at least on the animals that can be found closer to the shore by the whale watching boats. Whether this difference is due to the dissimilar oceanographic or geomorphologic characteristics of the archipelagos, or is the result of a seasonal prey shifting as it has been described in other species of the genus *Globicephala* (De Stephanis et al., 2008), should be clarified in further studies. In any case, the simple fact that the analysis of the AIS data has detected this subtle annual cycle proves the sensitivity of this methodology.

In addition to noise, the physical presence of boats may disrupt cetacean activity patterns, particularly when boats seek direct interactions (e.g. whale watching). In these cases, theoretical studies suggest that individuals often perceive boats as a risk, and therefore respond through avoidance and other anti-predatory tactics Pirotta et al. (2015). Cetaceans may begin to avoid particular areas if the disturbance reaches a certain threshold or if there is little cost to abandoning that location (Wright et al., 2011). It has also been observed that Marine mammals may temporarily move away during periods of heavy vessel activity but re-inhabit the same area when traffic is reduced (Bejder et al., 2006). Given the fact that the highest intensity in the whale watching activities in the Canaries happens during summer, one could expect that the whales would move far from the island (deeper waters) to avoid the vessels during this season. But the monthly mean depth of the whale watching events shows a clear yearly cycle were the animals slightly approach to the coast on summer and move to deeper waters in winter. This results seem to suggest a lack of avoidance in the long term behaviour of the short-finned pilot whales in the area. Nevertheless, this observation does not exclude more subtle avoidance effects, as the displacement of the more sensitive animals from the area of disturbance Bejder et al. (2006). Similarly, the whale watching effort is not homogeneously distributed among the optimal habitat of the short-finned pilot whales in the area. Hence, it could be also some habitat shift along the distribution area, changing the location of the animals, but not the depth. Despite this was not observed in the auto-correlation data of the whale watching positions registered in the study, subtle displacements at a constant depth would be difficult to detect using this methodology. Consequently, to accredit the presence or absence of avoidance effects, more comprehensive studies with individual identification of the cetaceans and its movements within the area of distribution would be necessary. If this avoidance reactions could be found, the concurrent determination of the whale watching effort using the proposed AIS methodology would allow to establish sustainable whale watching thresholds where avoidance does not occur.

Finally, the fine spatial resolution of whale watching effort obtained by this methodology is very promising. Not only as a tool to analyze the effects of different whale sighting intensities in future studies, but also to enforce the spatial regulation of the activities in a region. The existence of guidelines, regulations, or laws in an area is no guarantee of compliance with these guidelines (Parsons, 2012), the best guidelines can become inefficient if there is a chronic lack of enforcement. The most widespread method for effort regulation is to limit the number of licences, but this does not take into account the variable effort of each vessel either in time and space; neither the size, characteristic propeller noise, etc. The actual scientific methods to measure effort and behavioural effects like theodolite observations, could be useful to enforce regulations, but are either expensive or time consuming to cover big areas (Bejder et al., 2006; Schuler et al., 2019). The enforcement of exclusion (or limited effort) areas during sensitive seasons using the AIS-basead methodology will be very easy to track, making the identification of any vessel breaching the regulations a fully automated process. The methodology proposed here would be able to automatically distinguish when a vessel is just sailing through an exclusion zone, or when it is performing a whale watching activity in a prohibited or regulated area. The possibility to determine the position also allows to evidence concurrent vessels with the same group of whales, which could be useful for the enforcement too. Obviously, to achieve that, all the licensed ships in the region should be equipped with an AIS transponder. On the other hand, since this methodology can be used to record whale watching operations regardless of the vessel, it could be also used to detect and identify non-authorized ships performing whale sighting. To detect this illegal whale watching operations, not only the authorized vessels should have an AIS transponder, but all the touristic vessels operating in the area.

A public network of terrestrial stations would be essential to receive and archive high precision data from a large amount of vessels, as the ships sailing at 0-14 knots transmit AIS messages every 12 seconds, or every four seconds if they are changing course (Aarsæther and Moan, 2009). This high transmission rate, and the possibility to archive all the messages as open data, would allow more accurate calculation of the individual whale watching events that could be used for effort estimation, but also to regulation enforcement. This is specially interesting when the whale watching activities are performed in remote or difficult to access areas (Parsons, 2012). Considering that some of the fastest growing whale-watching industries are in developing countries, and is still an enormous potential for considerable growth in whale-watching operations in other developing nations (Parsons, 2012), the possibility to develop an automatic system to help the enforcement of regulations would be of great help in the future.

## 5 CONCLUSIONS

The proposed methodology to automatically estimate whale watching effort using AIS messages and EMODNET Digital Terrain Model has proven to be effective in two areas off the Canary Islands. The results obtained in this preliminary study are very encouraging, allowing to estimate seasonal and annual trends in the total amount of time that the cetaceans were exposed to whale watching activities. The results also provide detailed geographical distribution of the whale watching effort, and can be used to analyze subtle movements of the pods within their local distribution range.

As the proposed methodology relies heavily in the percentage of vessels equipped with AIS transponders, to achieve an optimum evaluation of the whale watching effort in a region, the competent authorities should promote its installation at least in all the ships authorized to perform whale watching activities. Ideally, any vessel capable of performing touristic activities in the area should have installed an AIS transponder to detect also illegal operations.

To effectively survey the whale watching area, a comprehensive study has to be performed in order to install enough AIS terrestrial stations to get a complete coverage and ensure the maximum reception of broadcasted messages. The system should be dimensioned considering the number of vessels to manage the simultaneous incoming messages. Once the system is operational, some level of open data policy should be established to grant transparent access to researchers and other stakeholders.

The methodology based on AIS messages has also proved to be successful in providing detailed spatial information about the whale watching effort. This characteristic is very promising to manage spatial whale watching regulations, specially to verify the enforcement of exclusion zones and areas with limited activity.

Having enough open data sets and making them available to science will contribute to the generation of knowledge and the creation of innovative products and services that have an impact not only on social well-being, but also on sustainability. This is the challenge for the competent administrations, which must ensure the relevance of this open data: more quality in the diversity of data and greater reflection to facilitate correlations with each other.

## AUTHOR CONTRIBUTIONS

JA conducted the analysis and led the project. FR conducted specific data analysis. All authors listed have made a substantial, direct and intellectual contribution to the work, and approved it for publication.

## ACKNOWLEDGMENTS

The authors wish to thank the Interreg MAC (2014-2020) Authority for the financial support and assistance to the MARCET II Project (MAC2/4.6c/392) and David Novillo for his assistance in gathering information from the whale watching pilots in Puerto Colón.

## DATA AVAILABILITY STATEMENT

The datasets generated for this study are available on request to the corresponding author.

